# Effects of altering histone post-translational modifications on mitotic chromosome structure and mechanics

**DOI:** 10.1101/423541

**Authors:** Ronald Biggs, Patrick Liu, Andrew D. Stephens, John F. Marko

**Affiliations:** Department of Molecular Biosciences, Northwestern University, Evanston, Illinois 60208; Department of Physics and Astronomy, Northwestern University, Evanston, Illinois 60208

**Author notes:** Correspondence to: John F. Marko: Department of Molecular Biosciences, Northwestern University, 2205 Tech Drive, Evanston, IL 60208-3500.

## Abstract

During cell division chromatin is compacted into mitotic chromosomes to aid faithful segregation of the genome between two daughter cells. Post-translational modifications (PTM) of histones alter compaction of interphase chromatin, but it remains poorly understood how these modifications affect mitotic chromosome stiffness and structure. Using micropipette-based force measurements and epigenetic drugs, we probed the influence of canonical histone PTMs that dictate interphase euchromatin (acetylation) and heterochromatin (methylation) on mitotic chromosome stiffness. By measuring chromosome doubling force (the force required to double chromosome length), we find that histone methylation, but not acetylation, contributes to mitotic structure and stiffness. We discuss our findings in the context of chromatin gel modeling of the large-scale organization of mitotic chromosomes.

## Introduction

Chromatin structure is important for many different cellular functions. A dramatic change in chromatin structure and organization occurs during the transition from interphase to mitosis as the open, diffuse, compartmentalized, and transcriptionally accessible interphase chromatin becomes compact, rod-like, and transcriptionally repressed in mitosis (Wang and Higgins, 2013; Doenecke, 2014; Oomen and Dekker, 2017). While most work studying mitotic chromatin rearrangement focuses on large chromatin-organizing complexes like cohesin, condensin, and topoisomerases (Vagnarelli, 2012), mitosis also is associated with characteristic changes to histone post-translational modifications (PTMs) (Wang and Higgins, 2013; Oomen and Dekker, 2017).

Histone PTMs are chemical changes to histones, typically to their tails, some of which are associated with different chromatin structures and densities (Rice and Allis, 2001; Wang and Higgins, 2013). Acetylation, notably of histone 3 lysine 9 (H3K9ac), is associated with euchromatin, which is loosely packed, gene rich, and actively transcribed (Doenecke, 2014) Methylation, notably H3K9me^3^ and H3K27me^3^, is associated with heterochromatin, which is densely packed and poorly transcribed (Rice and Allis, 2001; Wang and Higgins, 2013; Oomen and Dekker, 2017). Histone PTMs may also intrinsically alter chromatin packing by changing the charge of histones (acetylation) and introducing hydrophobic moieties to histones (methylation) (Rice and Allis, 2001; Doenecke, 2014). Recent cryo-EM data has shown that histones are often positioned such that histone tails can physically interact with other nearby histone tails (Bilokapic *et al.*, 2018), possibly enabling the alteration to chromatin structure.

Changes to histone PTMs are known to affect the structure and stiffness of cell nuclei during interphase. Increased euchromatin has been correlated with weaker nuclei (Chalut *et al.*, 2012; Krause *et al.*, 2013; Haase *et al.*, 2016) specifically decreasing the short-extension force response of nuclei, which is governed by chromatin stiffness, and contributes secondarily to long extensions (Stephens *et al.*, 2017). Chromatin stiffness also contributes to nuclear shape (Banigan *et al.*, 2017). Decreased chromatin-based nuclear rigidity caused by increased euchromatin has also been shown to cause abnormal nuclear morphology (Stephens *et al.*, 2018), which is an indicator of different cellular diseases, including cancers (Chow *et al.*, 2012). Increased heterochromatin has been shown to cause stiffer nuclei and resistance to abnormal nuclear morphology (Stephens *et al.*, 2017; Stephens *et al.*, 2018). Thus, the correlations between chromatin state and histone PTMs with nuclear stiffness and shape indicate underlying connections between histone PTMs and chromatin stiffness.

Some histone PTM changes are associated specifically with mitosis. Bookmarking is the process where some histone PTMs are retained or stabilized during mitosis, which is thought to preserve the cell’s transcriptional state through mitosis (Wang and Higgins, 2013; Doenecke, 2014; Oomen and Dekker, 2017). These marks are important for maintaining cellular identity and function. Several histone methyl marks, both euchromatic (*e.g.* H3K4me^3^) and heterochromatic (*e.g.* H3K9me^3^ and H3K27me^3^) are possibly increased or maintained in mitosis (Xu *et al.*, 2009; Park *et al.*, 2011). Increased H4K20me^1^ has also been associated with loading of condensin, which organizes chromatin in mitosis (Beck *et al.*, 2012). Another hallmark of mitosis is the dramatic reduction in overall histone acetylation (Park *et al.*, 2011; Zhiteneva *et al.*, 2017), which may be important for mitotic compaction or related to the lower transcriptional activity during mitosis (Wang and Higgins, 2013).

Histone PTMs may also intrinsically affect mitotic chromosome organization (Vagnarelli, 2012; Zhiteneva *et al.*, 2017). Recent experiments suggest that nucleosomes reconstituted using core histones from mitotic cells have a greater propensity to aggregate, compared to nucleosomes assembled using core histones from interphase cells (Zhiteneva *et al.*, 2017). This suggests that histone PTMs and their changes in mitosis may intrinsically affect mitotic compaction through nucleosome-nucleosome interactions. Other experiments have shown that DNA forms the underlying connectivity of mitotic chromosomes (Poirier and Marko, 2002; Sun *et al.*, 2011) and condensin in the central axis of mitotic chromosomes is discontiguous (Sun *et al.*, 2018; Walther *et al.*, 2018). While condensin provides the majority of the stiffness of mitotic chromosomes, it remains unclear how much chromatin-chromatin interactions could contribute to the stiffness of the mitotic chromosome.

To study the effects of altering histone PTMs on mitotic chromosome structure, we measured the doubling forces of captured mitotic chromosomes (Figure 1 and S1; the “doubling force” is the force required to double the length of a chromosome, and quantifies chromosome elastic stiffness in a chromosome-length-independent way). In order to test the hypothesis that alterations to histone PTMs affect the compaction of mitotic chromosomes, we studied the effects of the histone deacetylase inhibitors (HDACis), valproic acid (VPA) (Marchion *et al.*, 2005) and trichostatin A (TSA) (Yoshida *et al.*, 1990), on the levels of H3K9ac in mitosis and how they affect the stiffness of human mitotic chromosomes. We also tested how the histone demethylase inhibitor (HDMi), methylstat (MS), which is a Jumonji C-specific inhibitor (Luo *et al.*, 2011) (a key domain for several demethylases’ activity), alters the levels of H3K9me^2,3^ and H3K27me^3^ in mitosis, and affects the stiffness of human mitotic chromosomes. Our results show that HDACi treatments increase H3K9ac, but cause no change to the stiffness of mitotic chromosomes, while MS treatment increased canonical heterochromatin marks and the mechanical stiffness of mitotic chromosomes.

**Figure 1.**
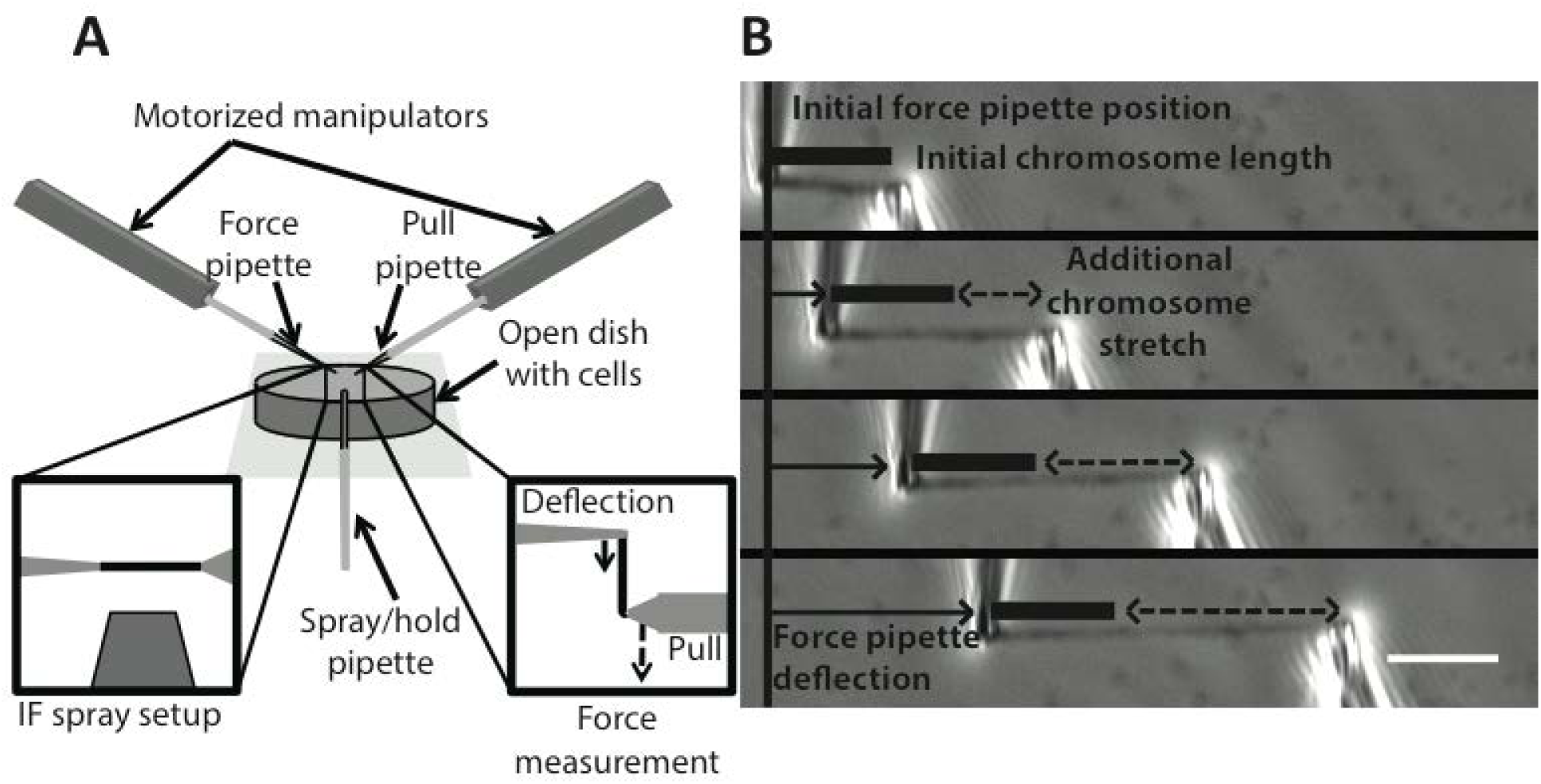
Experimental setup for chromosome micromanipulation, force measurement and image quantification. (**A**) Schematic of the single captured chromosome experimental setup. Single chromosomes were captured from mitotic HeLa cells in a custom-made well (Materials and Methods). Capture was performed after lysing the cell membranes with a PBS-Triton-X solution, where the chromosome was captured from the whole genome chromosome bundle (Figure S1). Once captured, the chromosome could be stretched for measurements of the doubling force or sprayed with fluorescent antibodies for immunostaining experiments. (**B**) An example of an experiment to measure the doubling force of a mitotic chromosome. The force (thin pipette on the left) and pull (larger pipette on the right) pipettes were aligned to be roughly perpendicular to the captured chromosomes. The pull pipette then moved away from the force pipette, stretching the chromosome (dashed line). The stretching of the chromosome would cause the force pipette to deflect (thin, rightward arrow) from its original position (thin, vertical line), which was used to calculate the force on the chromosome for the amount of stretch at that point. Chromosome initial length (thick bar) (measured by the distance from the center of the pipettes) and diameter (not shown) measured using a still image in ImageJ.

## Results

### HDACis increase H3K9ac on mitotic chromosomes but do not affect their stiffness

In order to investigate the role of histone PTMs on mitotic chromosome compaction, we studied the effects of histone hyperacetylation. We induced histone hyperacetylation using the histone deacetylase inhibitors (HDACi), valproic acid (VPA) and trichostatin A (TSA). Both VPA and TSA led to an increase in H3K9ac fluorescence intensity in fixed immunofluorescence (IF) (Figure S2A,B) and Western blots in interphase cells (Figure S2C). Having been able to induce hyperacetylation in interphase, we next tested whether the same treatment would cause histone hyperacetylation in mitosis. In fixed IF experiments of mitotic cells the average ratios of HDACi-treated to untreated H3K9ac acetylation levels were 1.4±0.1 for VPA and 2.3±0.3 for TSA (Figure 2A,B). In single captured chromosome experiments the average ratios of HDACi to untreated H3K9ac measurements were 1.8±0.2 for VPA and 2.3±0.6 TSA (Figure 2C,D). These results indicated that we were able to create hyperacetylated chromatin in mitosis.

**Figure 2.**
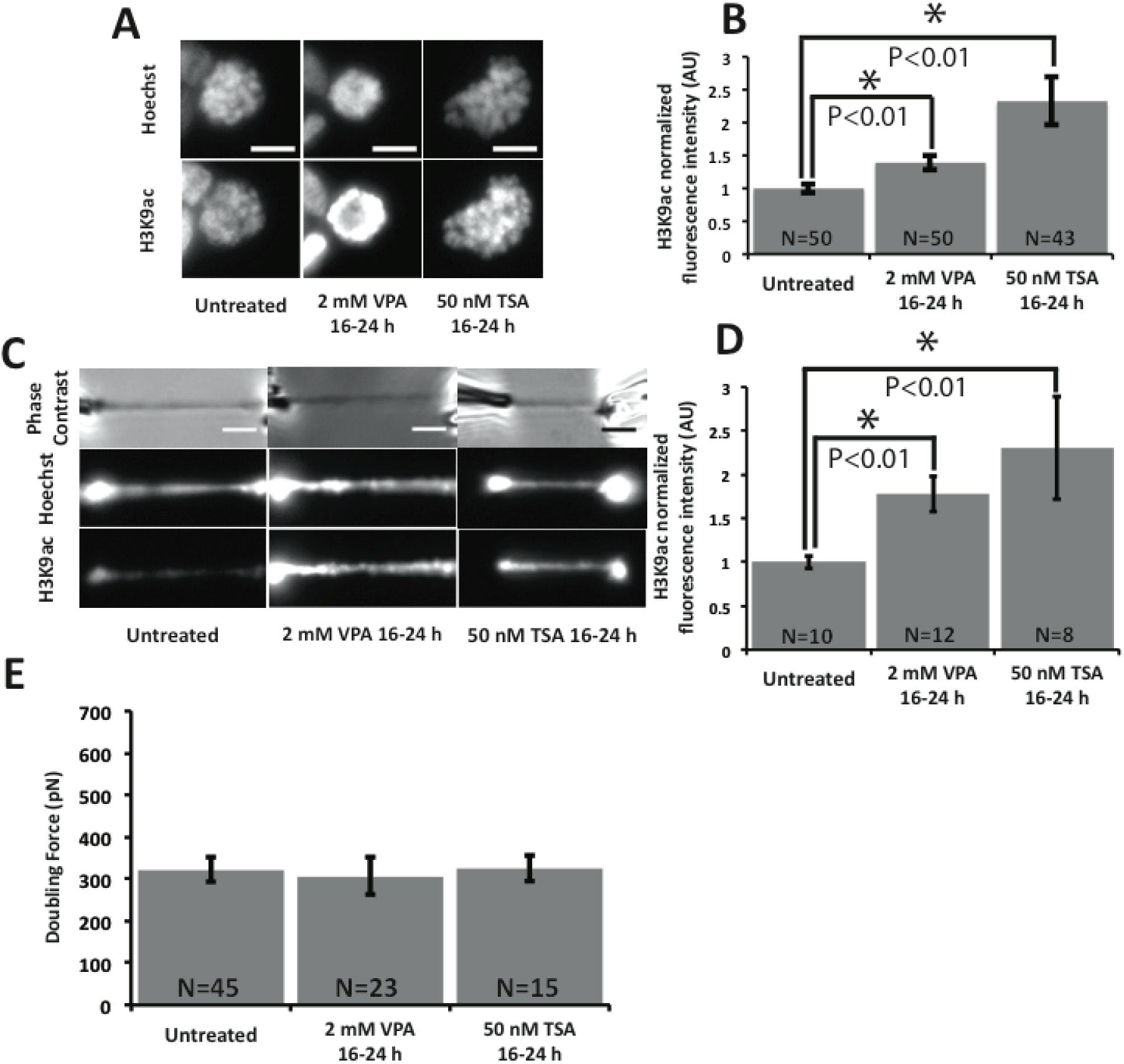
HDACis cause increased H3K9ac fluorescence in mitotic fixed cells and captured chromosomes, but have little effect on the stiffness of mitotic chromosomes. (**A**) Example representative images of levels of H3K9ac fluorescence measurement on fixed mitotic cells. Scale bar 10 μm. (**B**) Quantitative data of (**A**). The H3K9ac intensity ratio of untreated to 2 mM VPA 16-24 h treatment was 1.4±0.1 and is statistically significant. The H3K9ac intensity ratio of untreated to 50 nM TSA 16-24 h treatment was 2.3±0.3 and is statistically significant. (**C**) Example representative images of levels of H3K9ac fluorescence measurements on captured mitotic chromosomes. Scale bar 5 μm. (**D**) Quantitative data of (**C**). The H3K9ac intensity ratio of untreated to 2 mM VPA 16-24 h treatment was 1.8±0.2 and is statistically significant. The H3K9ac intensity ratio of untreated to 50 nM TSA 16-24 h treatment was 2.3±0.6 and is statistically significant. (**E**) Recorded doubling force for mitotic chromosomes from untreated and HDACi treated cells. The average chromosome doubling forces were 320±30 pN in untreated cells. The average doubling force was 310±40 pN in 2 mM VPA 16-24 h treated cells, statistically insignificant from untreated cells. The average doubling force was 330±30 pN in 50 nM TSA 16-24 h treated cells, statistically insignificant from untreated cells. Error bars represent standard error. Asterisk in bar graphs represent a statistically significant difference (*p* < 0.05). All *p* values calculated via *t* test.

Next we tested if this increase in acetylation would lead to a difference in stiffness for mitotic chromosomes, by measuring the doubling force of mitotic chromosomes extracted from untreated and HDACi-treated cells. Neither VPA nor TSA caused a statistically significant change in doubling force compared to untreated chromosomes (Figure 2E). The average chromosome doubling forces were 320±30 pN in untreated cells, 310±40 pN in VPA treated cells, and 330±30 pN in TSA treated cells. The lack of change was not due to changes of initial length or cross sectional area, as neither changed with HDACi treatments (Figure S2D,E).

Plotting the averaged doubling force against H3K9ac fluorescence for untreated and HDAC inhibited chromosomes, we found that there was no statistically significant correlation between H3K9ac measurements and doubling force in either untreated chromosomes or VPA treatments (Figure S2F). We do note that the TSA-treated chromosomes did show a statistically significant correlation between measured H3K9ac level and doubling force, with increasing acetylation leading to lower spring constant; however, when averaged over, there was no net effect of TSA treatment on chromosome spring constant. The correlation may be due to the specific mechanism of HDAC inhibition by TSA (no such correlation was observed for VPA), may reflect differences between specific chromosomes, or simply arise from the sample size being too small for this type of correlation analysis. Apart from this correlation, we concluded that hyperacetylation of histones through HDACi treatment does not affect the overall stiffness of mitotic chromosomes.

### Methylstat stiffens mitotic chromosomes and increases fixed cell histone methylation

Given that there was no overall effect of histone acetylation on chromosome doubling force, we wanted to test how altering histone methylation affects the stiffness of mitotic chromosomes. In order to induce hypermethylation, we used the histone demethylase inhibitor (HDMi) methylstat (MS), which increased both H3K9me^2,3^ and H3K27me^3^ as assayed via both Western blotting (Figure S3A,B) and fixed-cell IF in interphase cells (Figure S3C). Having been able to induce hypermethylation in interphase, we next tested whether the same treatment would cause histone hypermethylation in mitosis. In fixed IF experiments of mitotic cells the average ratio of MS to untreated H3K9me^2,3^ measurement was 1.6±0.1 while the average ratio of MS to untreated H3K27me^3^ measurement was 3.9±0.5 (Figure 3A,B). In contrast to the fixed IF experiments, MS did not cause a statistically significant change in H3K9me^2,3^ nor H3K27me^3^ measurement using antibodies microsprayed onto single captured chromosomes (Figure 3C,D). While unexpected, this data is explainable due to a lack of antibody accessibility and penetration into the more compact hypermethylated chromosomes, and the short antibody incubation time for our microspraying of captured chromosomes, relative to fixed IF staining (~10 min versus ~16 h).

**Figure 3.**
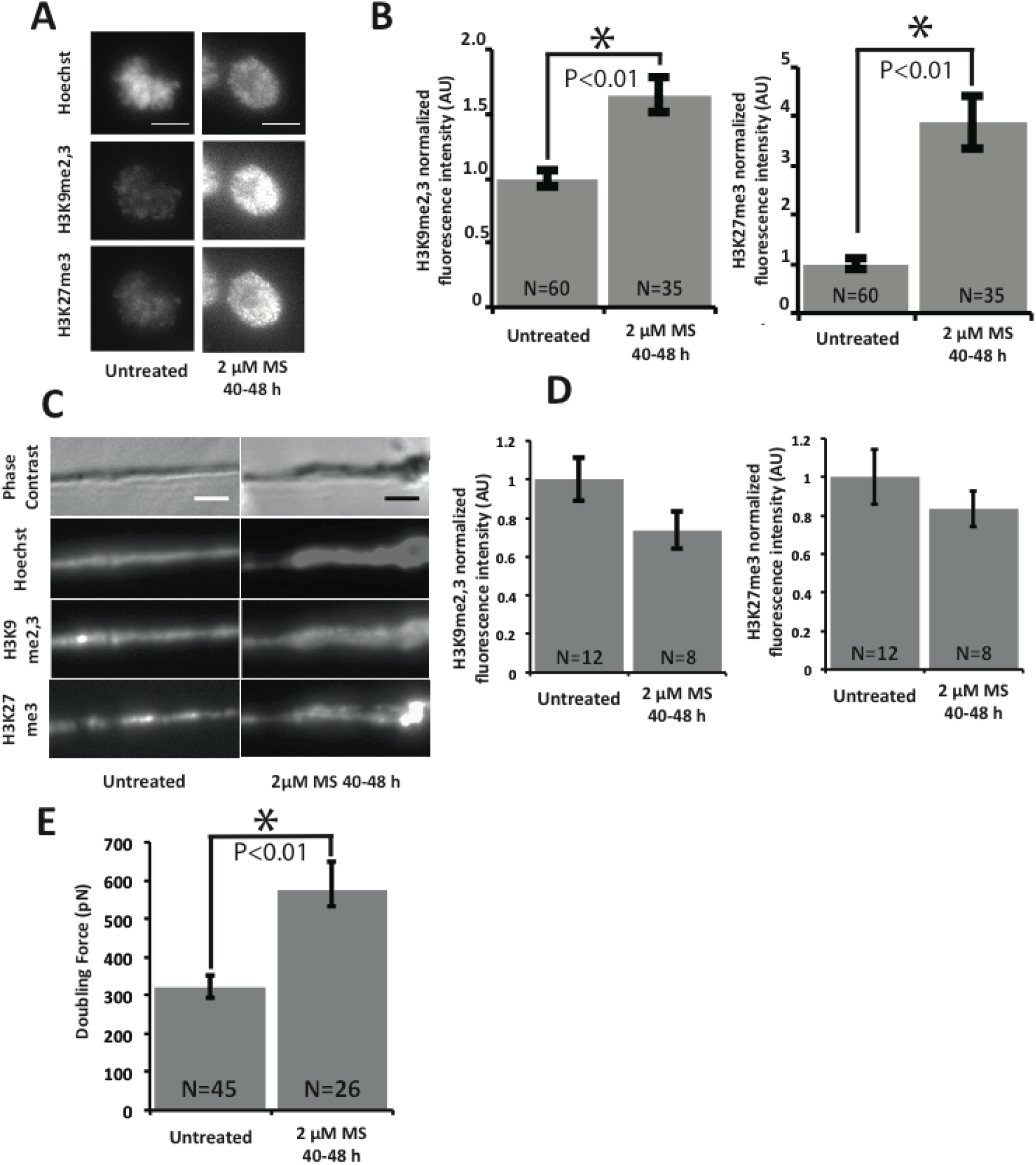
Methylstat (HDMi) treatment causes an increase in methylation for mitotic fixed cells and stiffens mitotic chromosomes. (**A**) Example representative images of levels of H3K9me^2,3^ and H3K27me^3^ fluorescence intensity on fixed mitotic cells. Scale bar 10 μm. (**B**) Quantitative data of (**A**). The H3K9me^2,3^ intensity ratio of untreated to 2 μM MS 40-48 h treatment was 1.9±0.1 and is statistically significant. The H3K27me^3^ intensity ratio of untreated to 2 μM MS 40-48 h treatment was 4.4±0.5 and is statistically significant. (**C**) Example representative images of levels of H3K9me^2,3^ and H3K27me^3^ fluorescence intensity on captured mitotic chromosomes. Scale bar 5 μm. (**D**) Quantitative data of (**C**). The H3K9me^2,3^ intensity ratio of untreated to 2 μM MS 40-48 h treatment was 0.73±0.10, statistically insignificant from untreated cells. The H3K27me^3^ intensity ratio of untreated to 2 μM MS 40-48 h treatment was 0.81±0.09, statistically insignificant from untreated cells. (**E**) Recorded doubling force for mitotic chromosomes from untreated and MS treated cells. The average chromosome doubling forces were 320±30 pN in untreated cells. The average doubling force was 580±40 pN in 2 μM MS 40-48 h treated cells, a statistically significant increase of ~%80 compared to untreated cells. Error bars represent standard error. Asterisk in bar graphs represent a statistically significant difference (*p* < 0.05). All *p* values were calculated via *t* test.

To determine if increased methylation caused mitotic chromosomes to become stiffer, we measured the doubling force of MS treated chromosomes. MS treatment caused a statistically significant increase of about 80% in the doubling force of mitotic chromosomes, consistent with more compact chromatin (Figure 3E). The average chromosome doubling forces were 320±30 pN in untreated cells and 580±40 pN in MS treated cells. This change was not due to a change in either the initial chromosome length or cross sectional area, as neither changed with MS treatment (Figure S3D,E).

Plotting doubling force against H3K9me^2,3^ measurements did not show any correlation in untreated or MS treated cells (Figure S3F left panels). Alternately, plotting doubling force against H3K27me^3^ measurements (in MS treated cells, but not untreated) suggests a potential correlation between H3K27me^3^ and chromosome stiffness (Figure S3F right panels). However, there may be limitations of antibody accessibility on the chromosomes, so this correlation must be regarded as preliminary at best. Our results do indicate that hypermethylation, via MS treatment, leads to robustly higher H3K27me^3^ levels, and causes chromosomes to become stiffer and possibly denser.

### Methylstat treatment does not change SMC2 levels

Since condensin is the most well known contributor to chromosome strength, we sought to check whether levels of condensin on mitotic chromosomes increased when treated with MS. Previous work has shown that chromosome stiffness is approximately linearly proportional to the amount of condensin on the chromosome (Sun *et al.*, 2018). We used antibodies against SMC2, a core subunit of condensin, to determine if there was a difference in fluorescence intensities between untreated and MS treated cells and captured chromosomes. The experiments did not show a difference as measured using fixed cellular immunofluorescence (Figure 4A,B) or for antibodies microsprayed onto captured chromosomes (Figure 4C,D), suggesting that the stiffening phenotype is independent of condensin loading.

**Figure 4.**
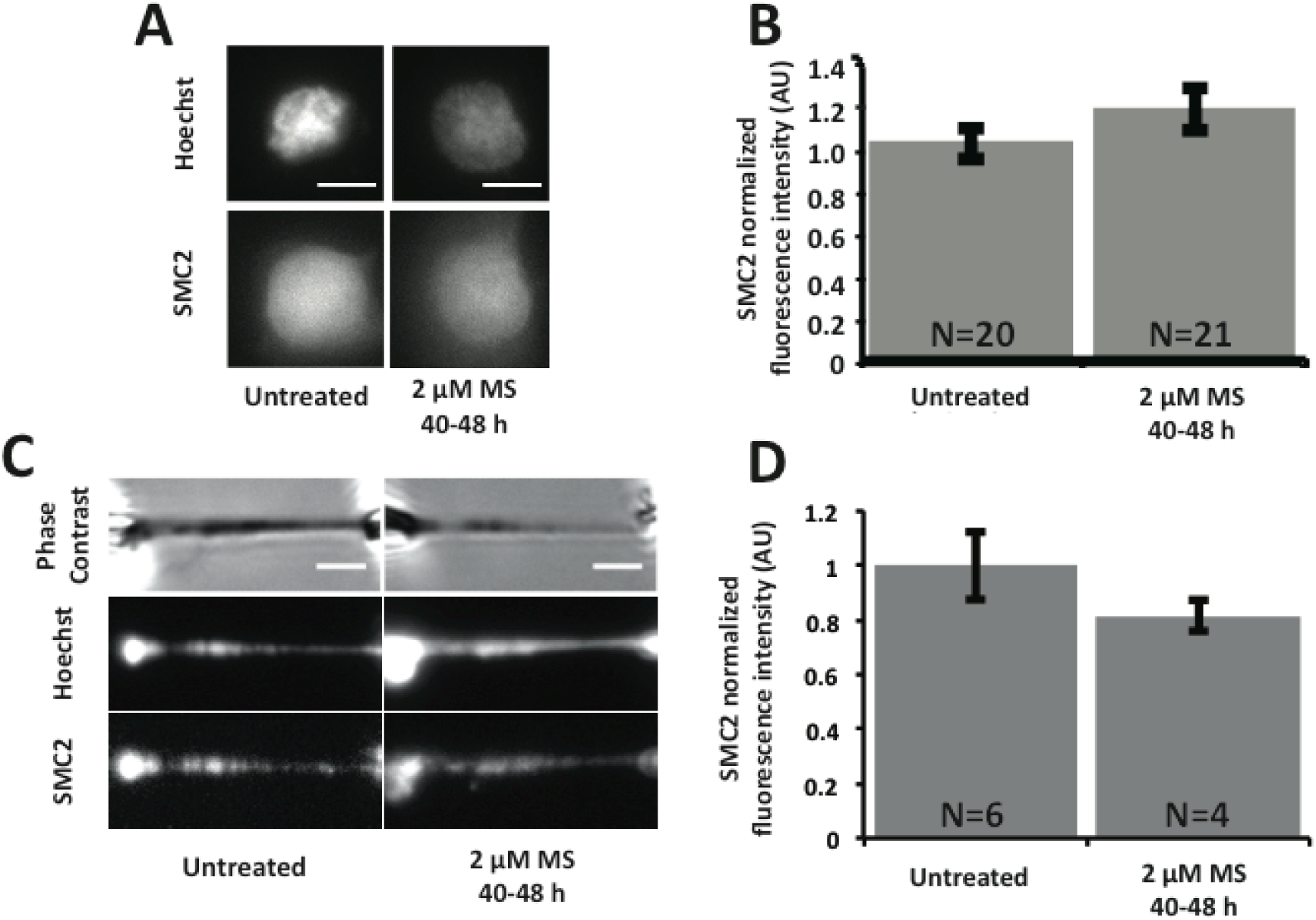
Methylstat treatment does not cause a change in SMC2 fluorescent levels. (**A**) Example representative images of levels of SMC2 fluorescence intensity on fixed mitotic cells. Scale bar is 10 μm. (**B**) Quantitative data of (**A**). The SMC2 intensity ratio of untreated to 2 μM MS 40-48 h treatment was 1.1±0.1, statistically insignificant from untreated cells. (**C**) Example representative images of levels of SMC2 fluorescence on captured mitotic chromosomes. Scale bar is 5 μm. (**D**) Quatitative data of (**C**). The SMC2 intensity ratio of untreated to 2 μM MS 40-48 h treatment was 0.82±0.05, statistically insignificant. All *p* values calculated via *t* test.

## Discussion

### Histone hypermethylation stiffens mitotic chromosomes, but hyperacetylation does not affect mitotic chromosome stiffness

Our data show that increasing histone acetylation (specifically H3K9ac level) by HDACi treatment does not affect chromosome stiffness in mitosis (Figure 2). Our original hypothesis had been that HDACi-induced histone hyperacetylation would weaken mitotic chromosomes. This hypothesis was based on the observations that histone acetylation is normally reduced in mitosis (Doenecke, 2014), and is thought to intrinsically affect nucleosome packing (Zhiteneva *et al.*, 2017). Furthermore, we expected to see weaker mitotic chromosomes since interphase hyperacetylated chromatin is decompacted (Doenecke, 2014) and hyperacetylating chromatin weakens the chromatin-dependent stiffness of interphase nuclei (Stephens *et al.*, 2017; Stephens *et al.*, 2018). However, our data indicate that mitotic chromosomes with hyperacetylated histones via HDACi treatment are overall just as stiff as mitotic chromosomes from untreated cells.

Unlike HDACi treatments, which do not change the doubling force of mitotic chromosomes, treatment by the HDMi MS causes increased histone methylation (assayed via H3K9 and H3K27 methylation) and a stiffer and likely denser mitotic chromosome without affecting SMC2 levels (Figures 3, 4). These results support our original hypothesis that the increase of histone methylation and propensity of mitotic histones to condense would stiffen mitotic chromosome as observed for interphase nuclei (Stephens *et al.*, 2017; Stephens *et al.*, 2018), but contrast with our results involving mitotic hyperacetylated histones. Our data indicate that this stiffening is not due to overloading of condensin, which suggests other mechanisms/complexes may affect chromosomal stiffness.

### Incorporating chromatin interactions into the model of mitotic chromosomes

To understand how chromatin may contribute to the overall stiffness of mitotic chromosomes, it is important to understand how mitotic chromosomes are organized. Early electron microscopy suggested that mitotic chromosomes are organized into loops of chromatin extending from a protein-rich core (Marsden and Laemmli, 1979). The currently heavily studied loop-extrusion model builds upon this classical bottlebrush model, describing how the bottlebrush is formed (Alipour and Marko, 2012; Goloborodko *et al.*, 2016; Gibcus *et al.*, 2018). In this model, chromatin loop-extruding complexes in the core of mitotic chromosomes create the bottlebrush structure. Non-histone chromatin-organizing complexes such as condensin and cohesin localize to the core of mitotic chromosomes and between sister chromatids, respectively (Ball and Yokomori, 2001; Piazza *et al.*, 2013), which according to the model function as loop-extruding enzymes. A broadly similar model of extruded chromatin loops organized by the protein complexes condensin and cohesin has been used to describe the vertebrate and yeast centromere as a chromatin spring (Ribeiro *et al.*, 2009; Stephens *et al.*, 2011; Lawrimore *et al.*, 2015).

We sought to incorporate the loop-extrusion model into the gel-network model of mitotic chromosomes. The gel-network model describes mitotic chromosomes as a gel of chromatin crosslinked by non-histone protein complexes, predominantly condensin (Figure 5A) (Poirier and Marko, 2002). There are two facets that govern the stiffness of a gel network: the density of crosslinks, and the pliability of the intervening cross-linked fibers (de Gennes, 1979). Older work has shown that condensin is responsible for about half of the spring constant of the kintetochore (Ribeiro *et al.*, 2009). Recent work has shown that condensin is approximately linearly correlated to the stiffness of mitotic chromosomes (Sun *et al.*, 2018), suggesting that most of the stiffness is governed by the chromatin loop-extruding elements, which are also apparently the primary crosslinking elements (Figure 5A). Previous work has shown that DNA/chromatin constitutes the underlying connectivity of mitotic chromosomes, which makes up the underlying fiber (Poirier and Marko, 2002; Sun *et al.*, 2011). These data also show that the loop-extruding proteins cannot form a contiguous core. In considering mitotic chromosomes as a gel, condensins comprise the major crosslinks while chromatin forms the underlying fiber. Both the lack of change in stiffness when histones are hyperacetylated and the lack of increase in condensin levels on hypermethylated histones suggests that perturbing histone PTMs does not affect the number of primary, condensin-based crosslinks.

**Figure 5.**
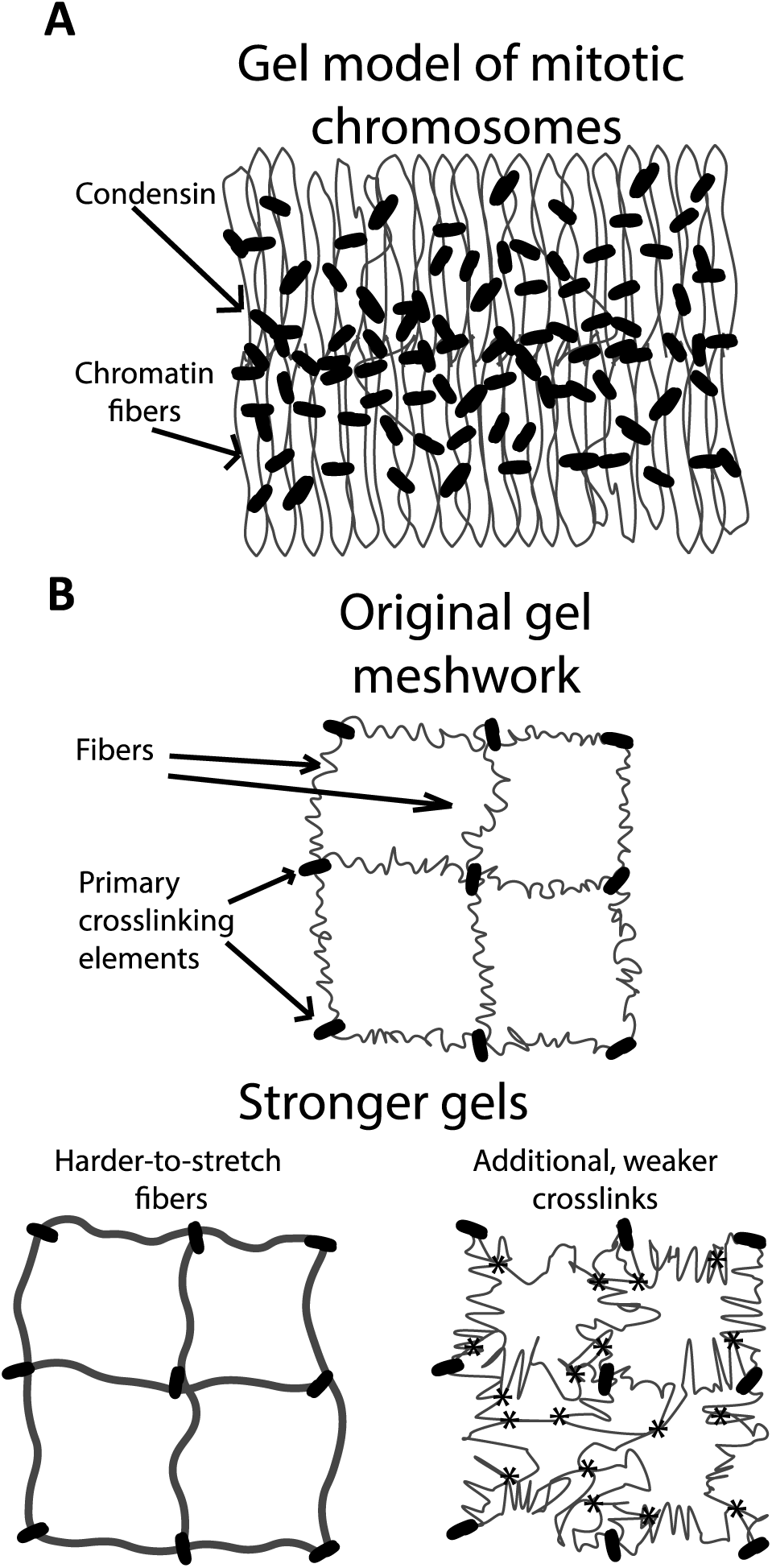
Model of mitotic chromosome. (**A**) Gel based model of mitotic chromosomes, demonstrating the crosslinking elements as condensin and the intervening fibers as chromatin. This model is compatible with different models of mitotic chromosomes including the loop-extrusion model, in which condensin can act both as a crosslinking element and the loop-extruding element. (**B**) Methods on which changes to the chromatin fiber or interactions of the chromatin fiber can stiffen a gel network. These models are not mutually exclusive and can be used to describe how increased histone methylation introduces an increase in stiffness to mitotic chromosomes. Neither of these effects are changed when histones are hyperacetylated in mitosis.

Since hyperacetylation of histones through HDACi treatments does not affect the stiffness of mitotic chromosomes, it cannot affect the amount of crosslinks or the ability of the chromatin fiber to be stretched. This is in contrast to interphase, where hyperacetylation weakens chromatin-based nuclear stiffness (Stephens *et al.*, 2017; Stephens *et al.*, 2018). This difference may be due to a lack of transcription in mitosis, acetyl-histone readers in mitosis, or other cell-cycle-dependent factors. These factors could actively decompact chromatin in interphase nuclei, but not in mitosis (Wang and Higgins, 2013; Doenecke, 2014). Furthermore, histone acetylation is drastically decreased in mitosis meaning that that the effect of increased histone acetylation via HDACi may be negligible for metaphase chromosomes. While a decrease in acetylation in mitosis coincides with a higher degree of compaction (Zhiteneva *et al.*, 2017), it appears that the increased acetylation of histones caused by our treatments with HDACis does not have an intrinsic effect on metaphase chromosome stiffness.

Our data suggest that hypermethylation of histones does affect mitotic chromosome structure, given the increased doubling force. Nucleosome-nucleosome interactions can stiffen mitotic chromosomes by either forming additional weaker crosslinks or the chromatin fibers themselves could become harder to stretch (Figure 5B). Neither of these hypotheses necessarily affect the primary crosslinkers, condensins. These two hypotheses are not mutually exclusive, although future experiments may be able to determine which of them is predominantly true. Further chromosome-manipulation experiments of the sort presented in this paper should be able to determine precisely which PTMs are responsible for the structural changes, as well as elucidate if the changes in chromosome mechanics we have observed are achieved by histones alone or if they require other proteins for their mediation.

A majority of work on the relation between histone PTMs and chromatin structure focuses on histone readers, but histone PTMs themselves may be intrinsically responsible for the stiffness change. It has been shown that chromatin reconstituted from mitotic histones aggregates more than chromatin reconstituted from interphase histones (Zhiteneva *et al.*, 2017). This analysis indicates that histone methylation is coupled to the structure and mechanics of mitotic chromosomes, in that a 3.4-fold increase in methylation is associated with an 80% increase in chromosome stiffness. This change in intrinsic condensation tendency may be facilitated by direct nucleosome-nucleosome interactions due to histone tails in the manner observed by (Bilokapic *et al.*, 2018). Our data do suggest that the potential increase of histone methylation, rather than decreased acetylation, contributes to tighter packing of nucleosomes during mitosis.

One must keep in mind that the metaphase chromosome, while organized as a chromatin gel, likely has an underlying radial-loop architecture, with an excess of condensin crosslinkers near the central chromatin “axes” (sketched in Fig. 5A). It is conceivable that weak, multivalent attractions between nucleosomes, such as those that might be mediated by methylated histone tails, could drive compaction of the denser axial region of metaphase chromatids without generating adhesion between the outer, less dense outer radial loop “halos”. Uncontrolled adhesion between nucleosomes must be avoided: once individual nucleosomes adhere to one another, the whole genome will stick together and form a droplet, a situation incompatible with chromosome segregation (Marko and Siggia, 1997). Multivalency could be a key ingredient, as it can permit a rapid “turn-on” of inter-nucleosome attraction with local nucleosome concentration, allowing the relatively weak loop-extrusion-compaction by condensins to compact the axial region sufficiently so that attractions turn on there, but not in the less dense loop halo. This scenario could explain how metaphase chromatids end up being dense in their axial interior while retaining mutually repulsive loop-halo exteriors, thus simultaneously achieving strong chromatin compaction while facilitating chromosome individualization and sister chromatid resolution, and also making the overall mechanics of metaphase chromosomes sensitive to additional nucleosome attractions associated with specific PTMs.

## Materials and Methods

### Cell culture and drug treatments

Human HeLa cells were maintained in DMEM (Corning) with 10% fetal bovine serum (FBS) (HyClone) and 1% 100x penicillin/streptomycin (Corning). The cells were incubated at 37°C and 5% CO_2_ for no more than 30 generations, and were passaged every 2-4 days. Experiments on captured chromosomes used cells that were allowed to recover 1-3 days before capture. Cells were freely cycling and not treated with drugs designed to affect or synchronize the cell cycle.

For epigenetic drug treatments, the cells were plated as above in drug-free DMEM and allowed to recover for ~8 h, then treated with 2 mM VPA (Sigma), 50 nM TSA (Sigma), or 2 µM MS (Cayman chemicals) all dissolved in DMEM. Chromosomes were then captured from the cells (see below) 16-24 h after treatment for VPA and TSA, or 40-48 h for MS treatments.

### Fixed immunofluorescence (IF)

Cells were grown on in small wells built on coverslips (Fisher) and treated as above. All solutions were diluted with and wash steps performed with PBS (Lonza) at room temperature, unless noted otherwise. Slides were washed, fixed in 4% paraformaldehyde (EMS), washed, permeabilized with 0.10-0.20% Triton-X 100 (USBio), incubated in 0.06% Tween 20 (Fisher), washed, and blocked in 10% goat serum (Sigma). The slides incubated with primary overnight at 4°C. The slides were then washed, incubated in secondary, incubated in Hoechst (Life Tech), washed and mounted.

Primary and secondary solutions were diluted in 10% goat serum. HDACi treatments were assayed using a 1:400 rabbit anti-H3K9ac (Cell Signaling 9649) primary and a 1:500 488-nm anti-rabbit IgG (Invitrogen A11034) secondary. HDMi treatments used a 1:100 mouse anti H3K9me^2,3^(Cell Signaling 5327) with a 1:1600 rabbit anti-H3K27me^3^ (Cell Signaling 9733) primary and a 1:500 of 488-nm anti-mouse IgG (Invitrogen A11001) with a 1:500 of 594-nm anti-rabbit IgG (Invitrogen A11037) secondary. Mitotic cells were identified by finding cells that showed compact mitotic chromosomes in the Hoechst channel. The final IF values reported are given by the fluorescence signal to background ratio of the antibody of interest over the Hoechst signal to background ratio. Averages and standard errors are divided by the average untreated values in normalized graphs.

### Single chromosome capture: setup and microscopy

Single chromosome capture experiments used an inverted microscope (IX-70; Olympus) with a 60x 1.42 NA oil immersion objective with a 1.5x magnification pullout at room temperature and atmospheric CO_2_ levels. Experiments were performed in less than 3 hours after removal from the incubator to ensure minimum damage to the cells being analyzed.

Prometaphase cells were identified by eye and lysed with 0.05% Triton-X 100 in PBS. All other pipettes were filled with PBS. After lysis, the bundle of chromosomes was held with a pipette. One end of a random, loose chromosome was grabbed by the force pipette (WPI TW100F-6), moved from the bundle and grabbed with the pulling pipette on the other end. The bundle was then removed to isolate the tracked and unbroken chromosome (Figure 1A and S1).

### Single chromosome capture: force measurement

An easily bendable force pipette and stiff pulling pipette were used for stretching chromosomes. Once captured, the pipettes were moved perpendicular to the chromosome, stretching the chromosome to roughly its native length. The stiff pipette was then moved 6 µm and returned to the starting position at a constant rate of 0.20 µm/sec in 0.04 µm steps using a LabVIEW program, while tracking the stiff and force pipette. Figure 1B shows an example stretch-deflection experiment. Deflection of the force pipette multiplied by its calibrated spring constant and divided by the distance between the pipettes (the stretch) was used to obtain the chromosome spring constant. Each chromosome was stretched at least 3 times to provide an accurate and reproducible measurement of the chromosome spring constant. The chromosome spring constant multiplied by its initial length gave the doubling force. The initial length was given by measuring the distance between the center of the pipettes in ImageJ and converting the pixels into microns while the chromosome was perpendicular to the pipettes. Chromosome cross sectional area was estimated as 0.25π*d*^2^ with chromosome diameter *d* calculated as the full width at half maximum of an ImageJ line scan.

### Single chromosome capture: immunofluorescence

After force measurements, the chromosome was lifted above the glass surface and microsprayed with a primary, secondary, and tertiary solution from a wide bore pipette, moving the chromosome between sprays. The solutions used 50 µL PBS, 36-38 µL H_2_O (Corning), 10 µL 5% casein (Sigma), and 2 µL each antibody. HDACi experiments used a rabbit anti-H3K9ac primary and a 488-nm anti-rabbit secondary. HDMi experiments used a mouse anti-H3K9me^2,3^ and a rabbit anti-H3K27me^3^ primary and a 488-nm anti-mouse IgG with a 594-nm anti-rabbit IgG secondary. The tertiary spray used Hoechst instead of an antibody.

### Western blots

Cells were grown in 100 mm dishes and treated as described in “cell culture and treatments”. TSA treatments were done at 200 nM. Cells were then harvested in PBS, centrifuged into a pellet, and lysed with RIPA buffer. The solution was then pelleted and the supernatant saved. The solution was then mixed with 2x Laemmli buffer, run on a 4-20% gradient SDS-PAGE gel, transferred to a nitrocellulose sheet, incubated in a primary solution, washed, and incubated in a secondary solution, then imaged.

### Statistics

For fixed immunofluorescence, the reported N refers to the number of technical replicates, *i.e.*, the total number of cells analyzed. The N measurements are furthermore from a set of biological replicates, *i.e.,* separate cell colonies on separate slides. All interphase-staining results are from data taken from two biological replicates. Mitotic-staining for H3K9ac and SMC2 were also obtained using 2 biological replicates. H3K9me^2,3^ and H3K27me^3^ data came from 4 biological replicates. For captured chromosomes, the reported N refers to each individual captured chromosome for both mechanical and immunofluorescence experiments; these experiments were from different slides (colonies) of cells and thus are independent biological replicates. Outliers were identified and discarded by using a generalized Studentized deviate test at α = 0.05. All p-values calculated using a T test. All averaged values are reported as average ± standard error.

## Acknowledgements

This work was supported by the NIH through grants R01-GM105847, U54-CA193419 (CR-PS-OC), ADS K99-GM123195, and a subcontract to grant U54-DK107980.

## Author contributions Statement

RB and JFM conceived and designed the research. RB, ADS, and PL conducted experiments. RB and PL analyzed data. RB, ADS and JFM wrote, read and approved the manuscript.

---

**Figure S1.**
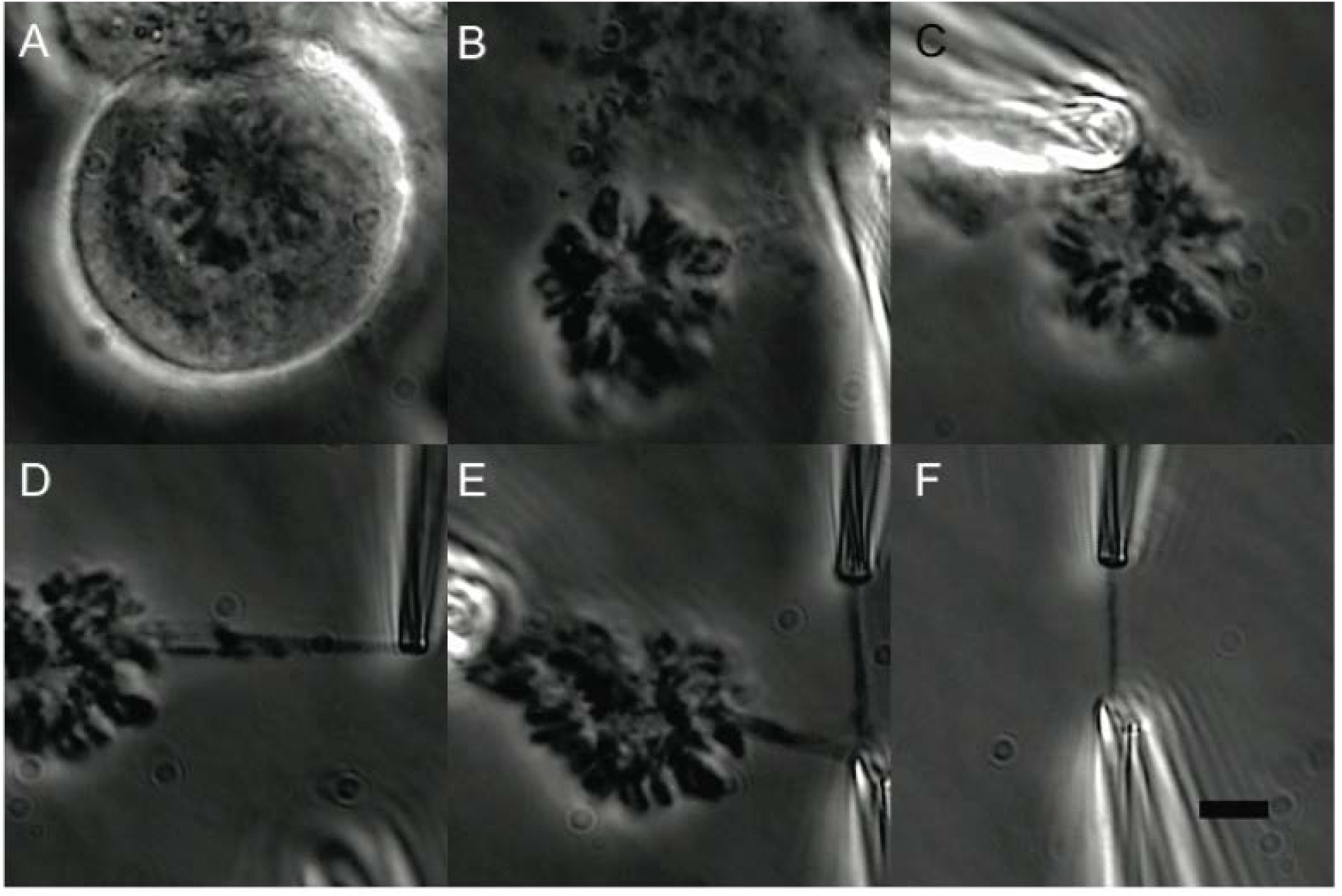
Example capture of single chromosome. (**A**) Morphology of a prometaphase mitotic HeLa cell: rounded morphology and clearly condensed chromosomes (**B**) The cell post Triton X-100 lysis (**C**) Chromosome bundle freed from the cell and after moving (**D**) The initial grab/aspiration of the chromosome into the force pipette (**E**) The second grab/aspiration of the other end of the chromosome into the stiff pipette (**F**) The chromosome after removal of the chromosome bundle. The chromosome is tracked as a single and unbroken object during the capture procedure. Scale bar is 5 μm.

**Figure S2.**
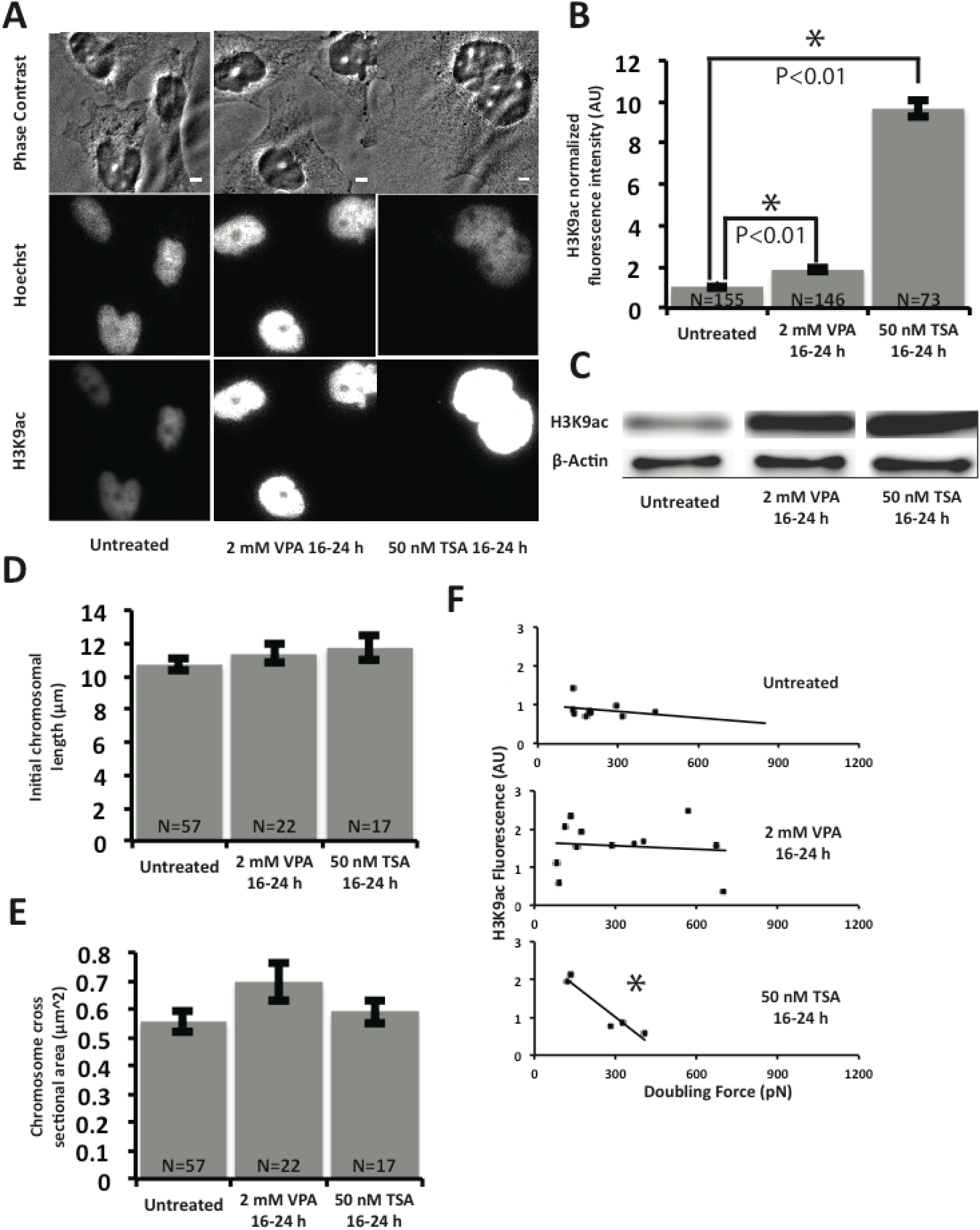
VPA and TSA treatment supplement. Both treatments cause hyperacetylation in interphase cells, but do not affect chromosomal initial length or cross sectional area. Only TSA displays a correlation between histone acetylation and doubling force. (**A**) Example representative images of levels of H3K9ac fluorescence measurement on fixed interphase cells. Scale bar is 5 μm. (**B**) Quantitative data of (**A**). The H3K9ac intensity ratio of untreated to 2 mM VPA 16-24 h treatment was 1.9±0.1 and is statistically significant.. The H3K9ac intensity ratio of untreated to 50 nM TSA 16-24 hr treatment was 9.7±0.1 and is statistically significant. (**C**) Western blot analysis of H3K9ac levels with β-Actin loading control in untreated, 2 mM VPA 16-24 h, and 50 nM TSA 16-24 hr treated cells. (D) Recorded initial length for mitotic chromosomes from untreated and HDACi treated cells. The average chromosome initial length was 10.7±0.3 μm in untreated cells. The average chromosome initial length was 11.4±0.6 μm in 2 mM VPA 16-24 h treated cells, statistically insignificant from untreated cells. The average chromosome initial length was 11.7±0.7 μm in 50 nM TSA 16-24 h treated cells, statistically insignificant from untreated cells. (**E**) Recorded cross sectional area for mitotic chromosomes from untreated and HDACi treated cells. The average chromosome cross sectional area was 0.56±0.04 μm^2^ in untreated cells. The average chromosome cross sectional area was 0.69±0.07 μm^2^ in 2 mM VPA 16-24 h treated cells, statistically insignificant from untreated cells. The average chromosome cross sectional area was 0.69±0.07 μm^2^ in 50 nM TSA 16-24 h treated cells, statistically insignificant from untreated cells. (**F**) Scatterplots of doubling force against H3K9ac fluorescence measurements. Using a linear fit the R^2^ were 0.06 for untreated, 0.01 2 mM VPA 16-24 h treatment, 0.91 for 50 nM TSA 16-24 h treatment. Error bars in SEM. All *p* values calculated via *t* test. All measurements recorded as statistically significant if *p* < 0.05. Asterisk in scatterplots represent a statistically significant correlation (*p* < 0.05).

**Figure S3.**
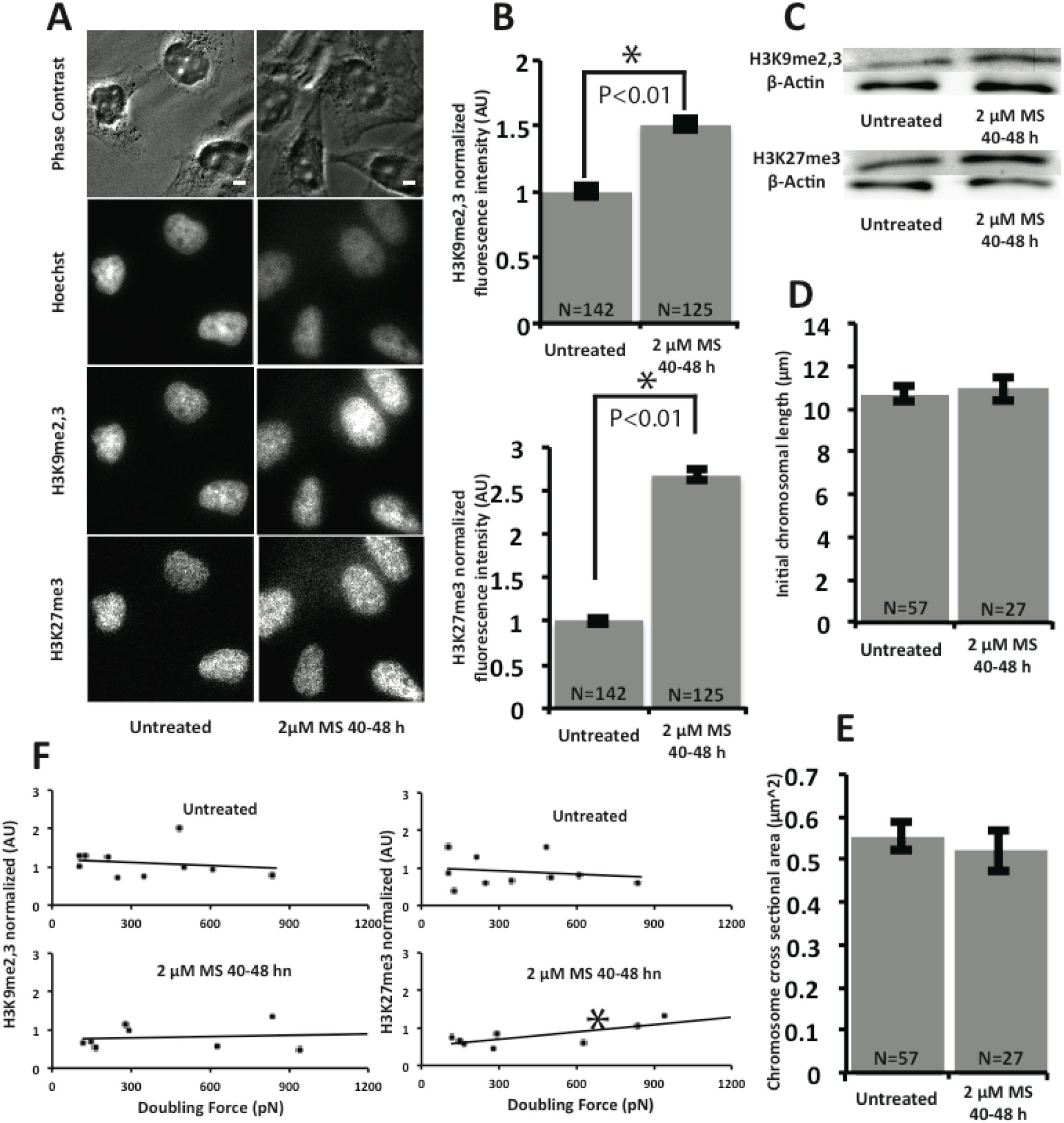
Methylstat treatment supplement. Treatment causes hypermethylation in interphase cells, but do not affect chromosomal initial length or cross sectional area. Only H3K27me^3^ fluorescence correlates with doubling force, only in methylstat treatment. (**A**) Example representative images of levels of H3K9me^2,3^ and H3K27me^3^ fluorescence measurement on fixed interphase cells. Scale bar is5 μm. (**B**) Quantitative data of (**A**). The H3K9me^2,3^ intensity ratio of untreated to 2 μM MS 40-48 h treatment was 1.5±0.1 and is statistically significant. The H3K27me^3^ intensity ratio of untreated to 2 μM MS 40-48 h treatment was 2.5±0.1 and is statistically significant. (**C**) Western blot analysis of H3K9me^2,3^ (top) and H3K27me^3^ levels with β-Actin loading control in untreated and2 μM MS 40-48 h treated cells. (D) Recorded initial length for mitotic chromosomes from untreated and MS treated cells. The average chromosome initial length was 10.7±0.3 μm in untreated cells. The average chromosome initial length was 11.0±0.6 μm in 2 μM MS 40-48 h treated cells, statistically insignificant from untreated cells. (**E**) Recorded cross sectional area for mitotic chromosomes from untreated and MS treated cells. The average chromosome cross sectional area was 0.56±0.04 μm^2^ in untreated cells. The average chromosome cross sectional area was 0.52±0.05 μm^2^ in 2 μM MS 40-48 h treated cells treated cells, statistically insignificant from untreated cells. (**F**) Scatterplots of doubling force against H3K9me^2,3^ and H3K27me^3^ fluorescence measurements. Using a linear fit the R^2^ were 0.03 for untreated H3K9me^2,3^, 0.03 for untreated H3K27me^3^, 0.01 for 2 μM MS 40-48 h treatment H3K9me^2,3^, 0.56 for 2 μM MS 40-48 h treatment H3K27me^3^. Error bars in SEM. All *p* values calculated via *t* test. All measurements recorded as statistically significant if *p* < 0.05. Asterisk in scatterplots represent a statistically significant correlation (*p* < 0.05).

## References

Alipour, E., and Marko, J.F. (2012). Self-organization of domain structures by DNA-loop-extruding enzymes. Nucleic Acids Res 40, 11202–11212.

Ball, A.R., Jr., and Yokomori, K. (2001). The structural maintenance of chromosomes (SMC) family of proteins in mammals. Chromosome Res 9, 85–96.

Banigan, E.J., Stephens, A.D., and Marko, J.F. (2017). Mechanics and Buckling of Biopolymeric Shells and Cell Nuclei. Biophys J 113, 1654–1663.

Beck, D.B., Oda, H., Shen, S.S., and Reinberg, D. (2012). PR-Set7 and H4K20me1: at the crossroads of genome integrity, cell cycle, chromosome condensation, and transcription. Genes Dev 26, 325–337.

Bilokapic, S., Strauss, M., and Halic, M. (2018). Cryo-EM of nucleosome core particle interactions in trans. Sci Rep 8, 7046.

Chalut, K.J., Hopfler, M., Lautenschlager, F., Boyde, L., Chan, C.J., Ekpenyong, A., Martinez-Arias, A., and Guck, J. (2012). Chromatin decondensation and nuclear softening accompany Nanog downregulation in embryonic stem cells. Biophys J 103, 2060–2070.

Chow, K.H., Factor, R.E., and Ullman, K.S. (2012). The nuclear envelope environment and its cancer connections. Nat Rev Cancer 12, 196–209.

Doenecke, D. (2014). Chromatin dynamics from S-phase to mitosis: contributions of histone modifications. Cell Tissue Res 356, 467–475.

de Gennes, P.G.d. (1979). Scaling concepts in polymer physics. Cornell University Press: Ithaca, N.Y.

Gibcus, J.H., Samejima, K., Goloborodko, A., Samejima, I., Naumova, N., Nuebler, J., Kanemaki, M.T., Xie, L., Paulson, J.R., Earnshaw, W.C., Mirny, L.A., and Dekker, J. (2018). A pathway for mitotic chromosome formation. Science 359.

Goloborodko, A., Imakaev, M.V., Marko, J.F., and Mirny, L. (2016). Compaction and segregation of sister chromatids via active loop extrusion. Elife 5.

Haase, K., Macadangdang, J.K., Edrington, C.H., Cuerrier, C.M., Hadjiantoniou, S., Harden, J.L., Skerjanc, I.S., and Pelling, A.E. (2016). Extracellular Forces Cause the Nucleus to Deform in a Highly Controlled Anisotropic Manner. Sci Rep 6, 21300.

Krause, M., Te Riet, J., and Wolf, K. (2013). Probing the compressibility of tumor cell nuclei by combined atomic force-confocal microscopy. Physical biology 10, 065002.

Lawrimore, J., Vasquez, P.A., Falvo, M.R., Taylor, R.M., 2nd, Vicci, L., Yeh, E., Forest, M.G., and Bloom, K. (2015). DNA loops generate intracentromere tension in mitosis. J Cell Biol 210, 553–564.

Luo, X., Liu, Y., Kubicek, S., Myllyharju, J., Tumber, A., Ng, S., Che, K.H., Podoll, J., Heightman, T.D., Oppermann, U., Schreiber, S.L., and Wang, X. (2011). A selective inhibitor and probe of the cellular functions of Jumonji C domain-containing histone demethylases. J Am Chem Soc 133, 9451–9456.

Marchion, D.C., Bicaku, E., Daud, A.I., Sullivan, D.M., and Munster, P.N. (2005). Valproic acid alters chromatin structure by regulation of chromatin modulation proteins. Cancer Res 65, 3815–3822.

Marko, J.F., and Siggia, E.D. (1997). Polymer models of meiotic and mitotic chromosomes. Mol Biol Cell 8, 2217–2231.

Marsden, M.P., and Laemmli, U.K. (1979). Metaphase chromosome structure: evidence for a radial loop model. Cell 17, 849–858.

Oomen, M.E., and Dekker, J. (2017). Epigenetic characteristics of the mitotic chromosome in 1D and 3D. Crit Rev Biochem Mol Biol 52, 185–204.

Park, J.A., Kim, A.J., Kang, Y., Jung, Y.J., Kim, H.K., and Kim, K.C. (2011). Deacetylation and methylation at histone H3 lysine 9 (H3K9) coordinate chromosome condensation during cell cycle progression. Mol Cells 31, 343–349.

Piazza, I., Haering, C.H., and Rutkowska, A. (2013). Condensin: crafting the chromosome landscape. Chromosoma 122, 175–190.

Poirier, M.G., and Marko, J.F. (2002). Mitotic chromosomes are chromatin networks without a mechanically contiguous protein scaffold. Proc Natl Acad Sci U S A 99, 15393–15397.

Ribeiro, S.A., Gatlin, J.C., Dong, Y., Joglekar, A., Cameron, L., Hudson, D.F., Farr, C.J., McEwen, B.F., Salmon, E.D., Earnshaw, W.C., and Vagnarelli, P. (2009). Condensin regulates the stiffness of vertebrate centromeres. Mol Biol Cell 20, 2371–2380.

Rice, J.C., and Allis, C.D. (2001). Histone methylation versus histone acetylation: new insights into epigenetic regulation. Curr Opin Cell Biol 13, 263–273.

Stephens, A.D., Banigan, E.J., Adam, S.A., Goldman, R.D., and Marko, J.F. (2017). Chromatin and lamin A determine two different mechanical response regimes of the cell nucleus. Mol Biol Cell 28, 1984–1996.

Stephens, A.D., Haase, J., Vicci, L., Taylor, R.M., 2nd, and Bloom, K. (2011). Cohesin, condensin, and the intramolecular centromere loop together generate the mitotic chromatin spring. J Cell Biol 193, 1167–1180.

Stephens, A.D., Liu, P.Z., Banigan, E.J., Almassalha, L.M., Backman, V., Adam, S.A., Goldman, R.D., and Marko, J.F. (2018). Chromatin histone modifications and rigidity affect nuclear morphology independent of lamins. Mol Biol Cell 29, 220–233.

Sun, M., Biggs, R., Hornick, J., and Marko, J.F. (2018). Condensin controls mitotic chromosome stiffness and stability without forming a structurally contiguous scaffold. Chromosome Res.

Sun, M., Kawamura, R., and Marko, J.F. (2011). Micromechanics of human mitotic chromosomes. Physical biology 8, 015003.

Vagnarelli, P. (2012). Mitotic chromosome condensation in vertebrates. Exp Cell Res 318, 1435–1441.

Walther, N., Hossain, M.J., Politi, A.Z., Koch, B., Kueblbeck, M., Odegard-Fougner, O., Lampe, M., and Ellenberg, J. (2018). A quantitative map of human Condensins provides new insights into mitotic chromosome architecture. J Cell Biol 217, 2309–2328.

Wang, F., and Higgins, J.M. (2013). Histone modifications and mitosis: countermarks, landmarks, and bookmarks. Trends Cell Biol 23, 175–184.

Xu, D., Bai, J., Duan, Q., Costa, M., and Dai, W. (2009). Covalent modifications of histones during mitosis and meiosis. Cell Cycle 8, 3688–3694.

Yoshida, M., Kijima, M., Akita, M., and Beppu, T. (1990). Potent and specific inhibition of mammalian histone deacetylase both in vivo and in vitro by trichostatin A. J Biol Chem 265, 17174–17179.

Zhiteneva, A., Bonfiglio, J.J., Makarov, A., Colby, T., Vagnarelli, P., Schirmer, E.C., Matic, I., and Earnshaw, W.C. (2017). Mitotic post-translational modifications of histones promote chromatin compaction in vitro. Open Biol 7.

